# Epicardial slices: a 3D organotypic model for the study of epicardial activation and differentiation

**DOI:** 10.1101/2020.08.07.219931

**Authors:** D. Maselli, R. S. Matos, R. D. Johnson, C. Chiappini, P. Camelliti, P. Campagnolo

## Abstract

The epicardium constitutes an untapped reservoir for cardiac regeneration. Upon myocardial injury, the adult epicardium re-activates the embryonic program, leading to epithelial-to-mesenchymal transition, migration and differentiation. Despite some successes in harnessing the epicardial therapeutic potential by thymosin β4 (Tβ4) pre-treatment, further translational advancements are hampered by the paucity of representative experimental models. Here we apply innovative protocols to obtain living 3D organotypic slices from porcine hearts, encompassing the epicardial/myocardial interface. In culture, our slices preserve the *in vivo* architecture and functionality, presenting a continuous epicardium overlaying a healthy and connected myocardium. Upon Tβ4 treatment of the slices, the epicardial cells become activated upregulating embryonic and EMT genes and invading the myocardium where they differentiate towards the mesenchymal lineage. Our 3D organotypic model enables to investigate the reparative potential of the adult epicardium, offering a new tool to explore *ex vivo* the complex 3D interactions occurring within the native heart environment.

## Introduction

The epicardium plays a crucial role during embryonic development of the heart (*1*), with lineage tracing studies indicating that the embryonic epicardium is a major source of cardiac fibroblasts (*2*), cardiac adipose tissue (*3*), and vascular smooth muscle cells and pericytes of the coronary vasculature (*4*).

In response to injury, such as myocardial infarction (MI), the adult epicardium re-activates the embryonic gene expression, including the transcription factors *Wilms’ tumor 1* (WT1) and *T box 18* (Tbx18) becoming activated (*5*). In this context, epicardial cells undergo epithelial-to-mesenchymal transition (EMT), before migrating into the myocardium and contributing to fibrosis, re-vascularization and cardiac repair through direct differentiation and paracrine stimulation (*5*–*7*).

The wound healing response resulting from epicardial activation is considerably increased by priming with thymosin-β4 (Tβ4) (*8*). Tβ4 is a 43 amino-acid long peptide that enhance the innate epicardial response following MI via epigenetic regulation of WT1 promoter (*9*) and drives epicardial EMT, ultimately increasing neovascularization and reducing pathological remodeling (*7, 9, 10*). In the context of post MI fibrosis, Tβ4 reduces collagen deposition both by attenuating profibrotic gene expression (*11*) and by resolving the immune response, as demonstrated *in vivo* in zebrafish and mouse models of cardiac injury and *in vitro* on human monocytes culture (*12*). However, the specific role of Tβ4 in fibroblast-mesenchymal differentiation from the epicardium has not been evaluated.

Given the limited understanding of the stimuli and mechanisms controlling the epicardial contribution to heart regeneration, new translational models are needed to investigate the role of endogenous and exogenous stimuli on the regulation of its reparative capacity.

Many studies describe epicardial activation as an evolutionarily conserved mechanism, but its efficiency varies greatly among different species: from the ability to regenerate significant portions of the adult heart in lower vertebrates, to a restricted reparative window during the early days after birth in small mammals (*13*). Despite some evidence of the newborn hearts’ recovery capacity (*14*), the role of the human adult epicardium in repair is unexplored due to the lack of a robust and representative model of epicardial function in large animals. Lineage tracing mouse models (*7, 15*) and primary or stem cell-derived human epicardial cells (*16*) yielded key findings, however a model combining the complex 3D *in vivo* environment and the relevance to human disease would provide further insights towards therapeutic applications. Myocardial slices were first described in the 1970s (*17*) and were subsequently used as a tool for electrophysiological and pharmacological studies (*18*). However, their applications were not extensively explored and widely recognized until the early 2000s, when their refinement provided major contributions to developing effective *ex vivo* models in cardiovascular research (*19*–*21*). The development of standardized protocols to obtain slices from a number of small and large mammals has allowed researchers to bridge the gap between *in vitro* and *in vivo* models, providing a valuable tool to study both the myocardial and non-myocardial cells of the heart (*19, 21*). In this study, we start from existing myocardial slice protocols to develop a reliable approach for the production and maintenance of epicardial slices from porcine hearts. We describe their use as a new, inexpensive and versatile *ex vivo* model to study the adult epicardium in large mammals. These epicardial slices include the epicardial/myocardial interface and maintain the typical 3D organization of cardiac tissue which enabled us to characterize the adult epicardium in large animal hearts. Through culturing the epicardial slices *ex vivo*, we recapitulate epicardial activation by Tβ4 treatment, which induces observable EMT, migration and differentiation of epicardial cells. Epicardial slice cultures combine the complexity of a 3D system with the flexibility of an *in vitro*/*ex vivo* system, providing a convenient research tool to dissect the role of the epicardium in cardiac homeostasis and repair and enabling further studies to harness the epicardial regenerative capacity for therapeutic purposes.

## Results

### 3D organotypic culture represents the epicardium/myocardium interface

We generated epicardial slices by developing an advanced embedding technique that protects the epicardial surface whilst achieving alignment of the tissue. Heart blocks are embedded in low melting agarose and flattened on to a compliant surface, to preserve epicardial viability. Using a high precision vibratome, we obtained a 400-500µm slice from the surface of the porcine heart. Structural analysis of histologically-processed slices showed that over 50% of the cells on the epicardium expressed the epicardial progenitor transcription factor WT1 (57.06±7.49% WT1^+^ epicardial cells, N of pigs=8, N of slices=8), with distinctive nuclear localization (Fig. 1A), while a small number of cells expressed the epithelial marker E-cadherin (Fig. 1B). Cells in the epicardium and the underlaying sub-epicardial layer also expressed the membrane marker Mesothelin (MSLN) (Fig. 1C). Immunostaining for α-sarcomeric actin (α-SA) and Connexin 43 indicated the retention of a structurally preserved and connected myocardium, presenting organized contractile units (Fig. 1, B and D) and intact gap junctions (Fig 1D). In addition, the slices presented numerous intact vascular structures characterized by the endothelial expression of CD31, including neuron-glial antigen 2^+^ (NG2) pericyte-covered capillaries and larger vessels surrounded by α-smooth muscle actin (α-SMA) -expressing smooth muscle cells (Fig. 1, D and F).

**Fig. 1.**
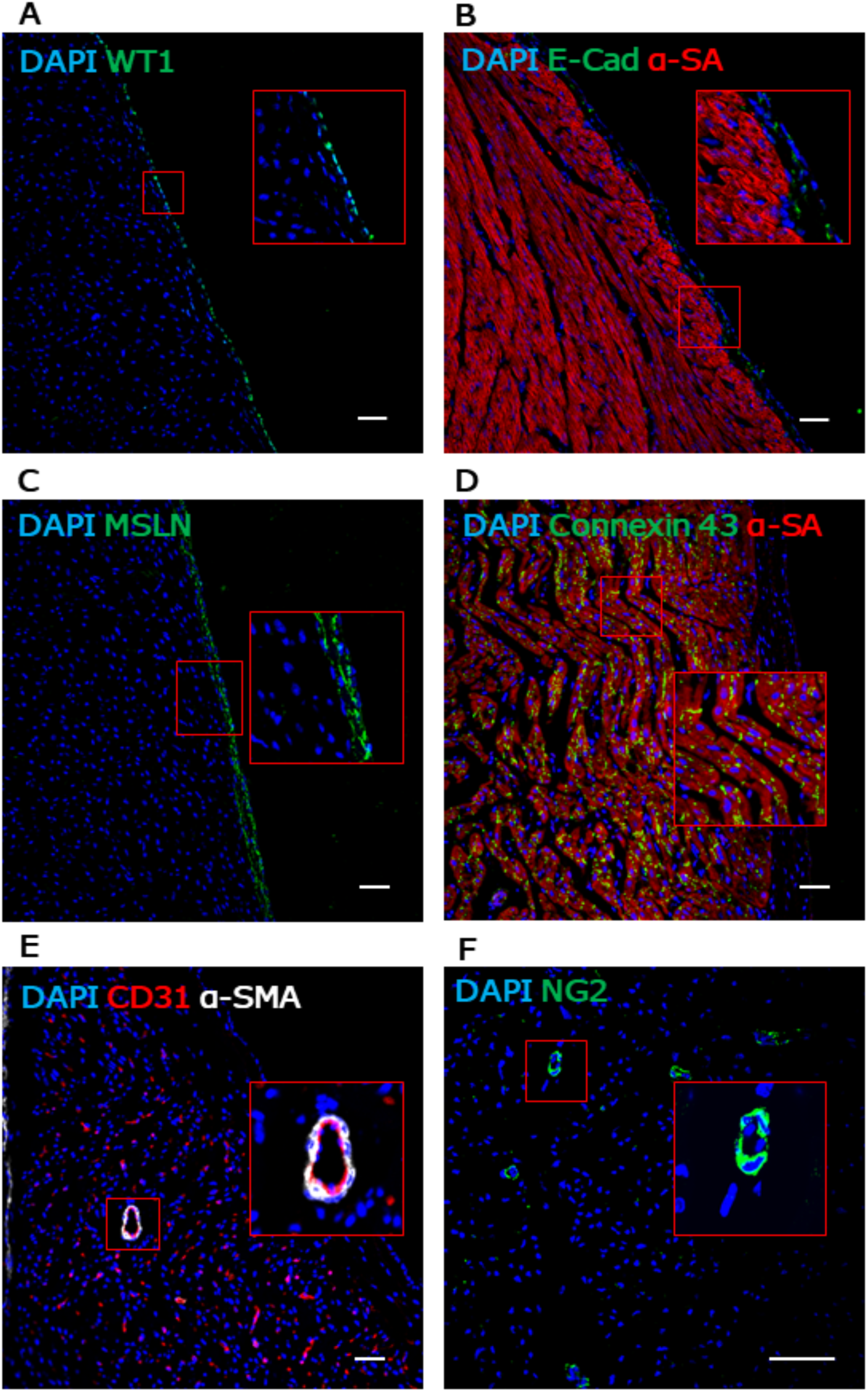
Porcine epicardial slices retain native morphology. Confocal microscopy analysis of formaldehyde-fixed OCT-embedded porcine epicardial/myocardial slices confirms the presence of an intact epicardial layer identified by the expression of WT1 (A) the sporadic expression of E-cadherin (B) and extensive positivity to mesothelin (MSLN) (C). The cardiomyocytes within the slice display intact α-sarcomeric actin structures (B) and gap junctions connexin 43 (D), indicative of well-preserved myocardial architecture. CD31 identifies vascular structures (E), including arterioles surrounded by smooth muscle cells expressing α-SMA (E) and capillaries associated with the occasional NG2^+^ pericytes (F). Nuclei are labelled with DAPI. Scale bar, 50µm.

Freshly cut slices presented an undamaged epicardial monolayer, covering the underlying connective and myocardial tissues. Therefore, this *ex vivo* model maintains the structural and cellular composition of the intact heart tissue and can be used to explore the epicardial/myocardial interface *ex vivo*.

### Culture maintains the integrity of epicardial slices

We developed two different culture systems to allow *ex vivo* culture of epicardial slices: a simpler and inexpensive ‘static’ method and a more controlled and tunable ‘dynamic’ method. The static method advanced the most recent protocols for the air-liquid interface culture of myocardial slices (*21*). The dynamic method developed a perfusion bioreactor system (flow rate 4ml/min) equipped with a feedback loop control system capable of the real time monitoring and adjustment in the culture medium of the dissolved oxygen level at 21% and the pH at 7.4. We compared the static and dynamic systems by measuring cell viability, apoptosis and proliferation rate after 24 and 48 hours. Calcein acetoxymethyl (Calcein AM) staining was used to quantify the live/metabolically active cells on the surface of fresh slices. All slices retained the characteristic cobblestone-like morphology of the epicardium, confirming the presence of undamaged epicardial cells up to 48h (Fig. 2A). Confocal microscopy quantification of fluorescence showed that both static and dynamic cultures maintained the initial slice viability over a period of 48h, with a remarkable viable area of over 50% throughout (Fig. 2B). Histological quantification (Fig. 2C) revealed that freshly cut slices presented minimal levels of apoptosis, with 1.57±0.41% of dying cells in the whole slice (Fig. 2D), although the epicardium displayed a slightly higher percentage of apoptotic cells (7.42±1.69%, Fig. 2E) compared to the myocardium (1.03±0.32%, Fig2F). The level of apoptosis increased when cultured in static conditions for 48h, especially in the epicardium (24h 37.09±9.06%, vs. T0 p=0.0005; 48h 31.24±4.54%, vs. T0 p=0.0061, Fig. 2E), whilst the myocardium displayed a lower level and delayed onset of apoptosis, starting at 48h of culture (48h 11.08±3.88%, vs. T0 p=0.002, Fig. 2F). Slices cultured in the dynamic system instead underwent only a moderate and non-significant increase in the apoptosis within the first 24h, which did not worsen after 48h (Fig. 2, D to F). Proliferating cell nuclear antigen (PCNA) staining in fresh slices, showed low basal proliferation levels for both the epicardium and myocardium (Epicardium 0.14±0.08%; Myocardium 0.20±0.04%, Fig. 2, G to J), indicating an overall quiescent physiological state. This basal proliferation rate was maintained in static cultures, whilst the dynamic culture system stimulated a robust proliferation of epicardial cells which was significantly increased at 48h of culture when compared to static cultures (Dynamic 48h 8.00±3.83% vs. Stat 48h p=0.0391, Fig. 2, H to J). Indeed, slice generation and culturing did not upregulate the expression of embryonic epicardial transcription factors *WT1, Tbx18* and *Transcription factor 21* (*TCF21*), independently of the culture protocol (Fig. 2, K to M).

**Fig. 2.**
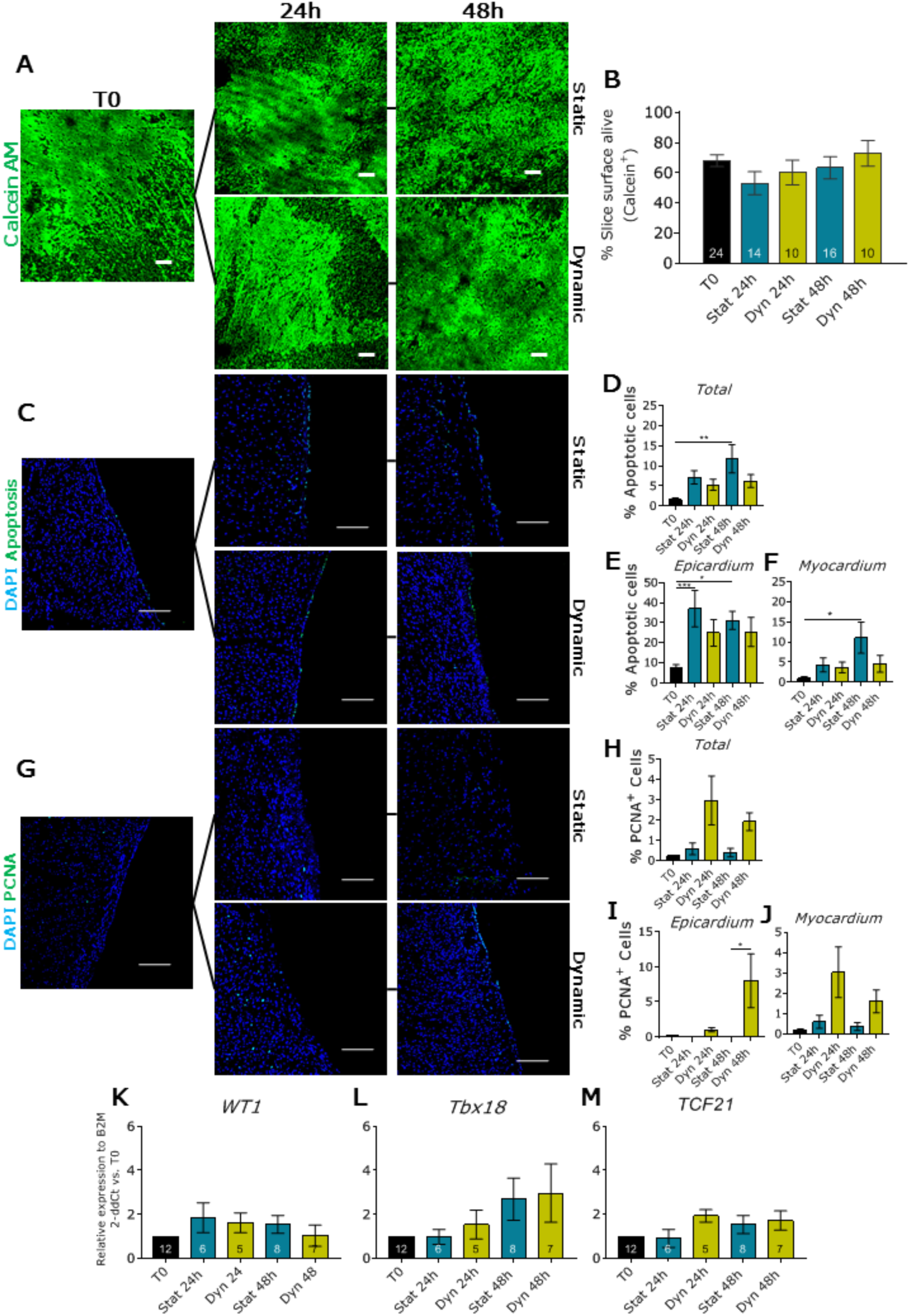
Culture preserve slice viability. Cell viability in fresh cut slices (T0) and after 24h and 48h of static (Stat) and dynamic (Dyn) culture (A and B). The epicardium maintains its morphology (A) and viability (B) upon culture, as quantified by average percentage of calcein AM positive area. N of Pigs = 6-18, number of slices displayed in graph. Apoptotic cells evaluated by ApopTag detection kit assay (C to F). Representative confocal images (C) and apoptotic cell fraction as a function of culture time and conditions in the entire slice (D), in the epicardial (E) and myocardial area (F). N of pigs = 3-6, number of slices: T0=13, Stat 24h=7, Dyn 24h=4, Stat 48h=7, Dyn 48h=5. Cell proliferation rate evaluated by PCNA antigen staining (G to J). Representative confocal images (G) and proliferating cell fraction in the whole slice (H), the epicardial (I) and the myocardial area (J). N of pigs = 5-6, number of slices: T0=5, Stat 24h=7, Dyn 24h=7, Stat 48h=5, Dyn 48h=7. Relative gene expression in the whole slices evaluated qPCR (K to M): expression of *WT1* (K), *Tbx18* (L), and *TCF21* (M) relative to basal expression at T0. N of Pigs = 3-7, number of slices displayed in graph. All graphs display data as mean with SEM. *=p ≤ 0.05, **=p ≤ 0.01, ***=p ≤ 0.001. Scale bar, 100µm.

Taken together, these results indicated that static culture conditions lead to moderate levels of apoptosis and no proliferation in epicardial slices, whilst the constant oxygenation and circulation of nutrients, as well as strict control of the pH afforded by the dynamic system, stimulated epicardial cell proliferation without activating embryonic epicardial genes and protected the myocardium from apoptosis.

### Tβ4 preserves epicardial viability and promotes activation and EMT

As the static culture system had a limited effect on epicardial cell proliferation, we decided to use it to assess the effect of the cardioprotective peptide thymosin β4 (Tβ4) (*22*) on the *ex vivo* epicardial tissue. Calcein AM staining showed increased viability of Tβ4 treated slices at 48h compared untreated cultures (Static 48h 62.32± 6.70% vs. Tβ4 48h 88.68± 2.57%, p=0.0349, Fig. 3, A and B). While the average apoptosis in the whole slice remained unaffected by Tβ4 treatment (Fig. 3, C and D), in the epicardium Tβ4 protected from the statistically significant increase in cell death observed in untreated cultures (Fig. 3E). This effect was unique to the epicardium, as the myocardium apoptosis increased independently of Tβ4 (Myocardium: Tβ4 48h 16.37±6.02%, vs. T0 p=0.0011, Static 48h 11.79± 3.78%, vs. T0 p=0.0022, Fig. 3F). Similarly, cell proliferation remained unchanged in the whole slice (Fig. 3, G and H), showing a significant and isolated increase in the epicardium (Fig. 3I), and no differences in the myocardium (Fig. 3J). Tβ4 treatment upregulated 3 to 6-fold the expression of epicardial transcription factors WT1, Tbx18 and TCF21 as compared to the untreated control slices (Fig. 4, A to C). For comparison, the expression of these transcription factors in myocardial slices was barely detectable levels and neither culture or Tβ4 supplementation resulted in significant upregulation (*data not shown*). We also observed a strong overexpression of Snails transcription factors (*Snai1* and *Snai2*) (Fig. 4, D and E) and *twist family bHLH transcription factor 1* (*TWIST-1*) (11-fold at 24h and 18-fold at 48h, Fig. 4F), which suggests Tβ4-dependent activation of the EMT pathways.

**Fig. 3.**
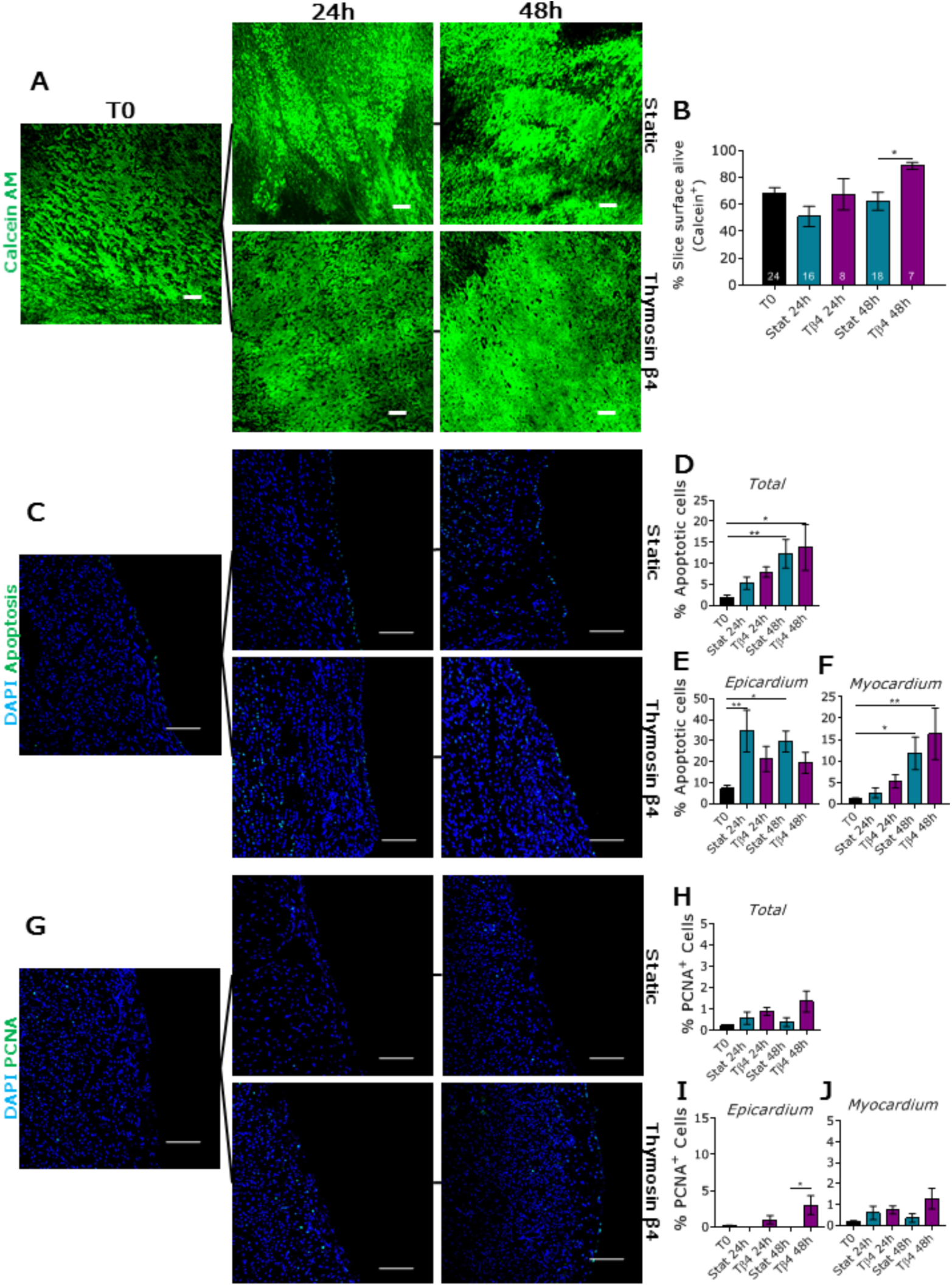
Tβ4 treatment alters epicardial cell viability and proliferation. Cell viability in fresh cut slices (T0), after 24h and 48h of control static (Stat) culture and thymosin β4 (Tβ4) treatment (A and B). Representative confocal images (A) and quantification (B) indicates an increase in the percentage of live area upon Tβ4 treatment at 48h. N of Pigs = 4-14, number of slices displayed in graph. Apoptotic cells quantification using ApopTag detection kit (C to F), representative images (C) and graphs indicating apoptotic cells as a function of culturing time and conditions: throughout the entire slice (D), in the epicardial (E) and the myocardial area (F). N of pigs = 3-6, number of slices: T0=13, Stat 24h=7, Tβ4 24h=5, Stat 48h=7, Tβ4 48h=5. Percentage of proliferative cells assessed by PCNA staining (G to J). Representative images of the PCNA expression among the different cultures (G). Quantification of the percentage of proliferative cells normalized on DAPI nuclear staining, in the whole slice (H), in epicardial (I) and myocardial area J). N of pigs = 5-6, number of slices: T0=5, Stat 24h=5, Tβ4 24h =6, Stat 48h=5, Tβ4 48h=8. All graphs display data as mean with SEM. *=p ≤ 0.05, **=p ≤ 0.01. Scale bar, 100µm.

**Fig. 4.**
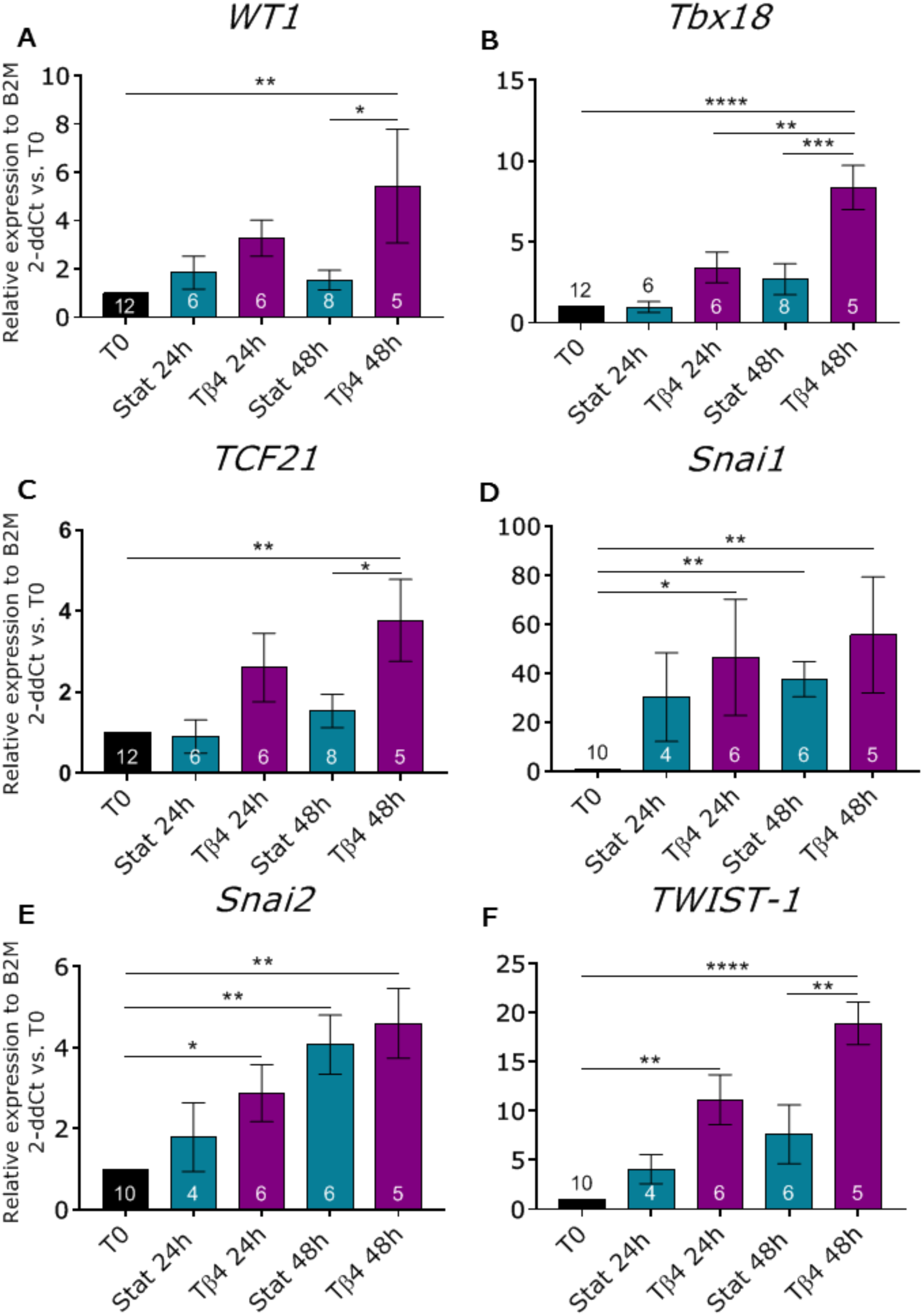
Tβ4 stimulates epicardial embryonic gene expression and EMT in the slices. Gene expression analysis showing relative expression of epicardial transcription factors *WT1* (A), *Tbx18* (B) and *TCF21* (C) and EMT markers *Snai1* (D), *Snai2* (E) and *TWIST-1* (F), as compared to freshly cut slices (T0). N of Pigs = 2-7, number of slices displayed in graphs. All graphs display data as mean with SEM. *=p ≤ 0.05, **=p ≤ 0.01, ***=p ≤ 0.001, ****p ≤ 0.0001.

Tβ4 treatment protected the viability of the epicardium on the slice and promoted epicardial cell proliferation. Moreover, Tβ4-treatment enhanced the expression of genes related with epicardial activation and EMT differentiation.

### Tβ4 changes WT1^+^ cells distribution and induces differentiation

Activation of epicardial cells increase their migratory capacity and differentiation potential (*8*). To estimate the re-activation and migration of epicardial cells in our *ex vivo* model we measured the localization of WT1^+^ cells in respect to the epicardial surface. In freshly prepared slices, WT1^+^ cells were largely confined to the surface of the slice, in the epicardial layer (T0: 0-50µm 91.51±3.77%, >50µm 8.48±3.77%, Fig. 5, A and B). Importantly, static culture did not change this distribution, whilst Tβ4 treatment progressively decreased the number of WT1^+^ cells in the epicardial layer starting at 24h (0-50µm: Tβ4 24h 81.24±5.03% vs. T0, p=0.0089), with a further reduction at 48h (0-50µm: Tβ4 48h 71.42±7.85% vs. T0 p=0.0129, Fig. 5, A and B). As a result, the percentage of WT1^+^ cells found in the subepicardial space and underlying myocardium increased in a time-dependent fashion (>50µm: Tβ4 24h 18.35±5.03% vs. T0, p=0.0089; Tβ4 48h 28.58±7.85 vs. T0, p= 0.0129, Fig. 5, A and B). Importantly, the total number of WT1^+^ cells in the whole slice remained overall unchanged during culture and among treatments despite a small but significant increase in proliferation within the epicardium, indicating that the redistribution of WT1^+^ cells after Tβ4 treatment might result from an increase in cell motility (Fig. 5C).

**Fig. 5.**
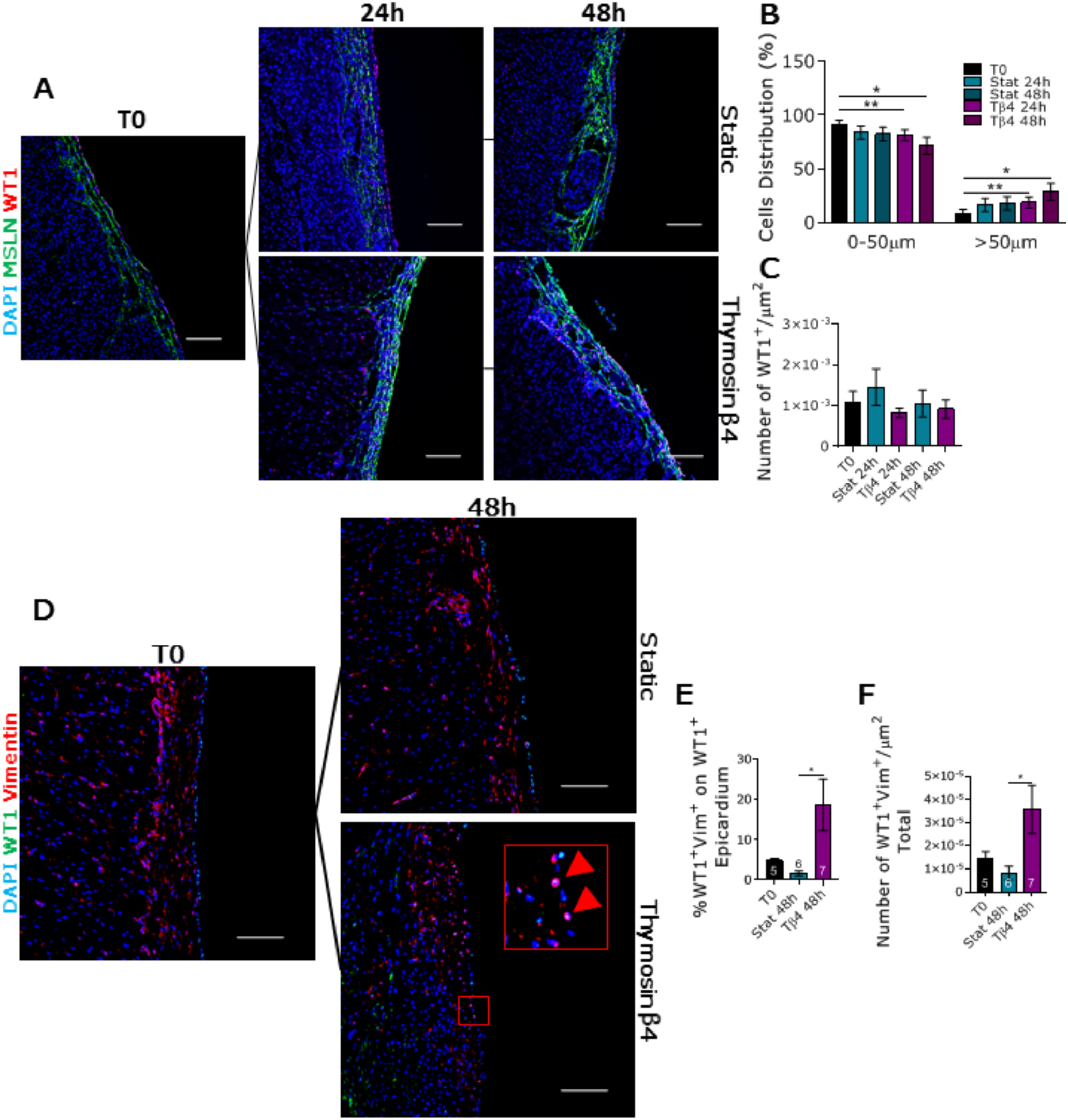
Tβ4 stimulates WT1^+^ epicardial cell migration and differentiation. WT1^+^ epicardial cells distribution evaluated as average distance from the epicardial monolayer. Representative images of the distribution of WT1 and MSLN expression at different time points (A). Graph indicating the percentage of WT1^+^ cells retained in the epicardial layer (0-50µm) or present in the myocardial area (>50µm) in different culture conditions (B). Graph indicating the quantification of WT1^+^ cells in the whole slices (C). N of Pigs = 5-7, number of slices: T0=8, Stat 24h=8, Tβ4 24h=8, Stat 48h=7, Tβ4 48h=5. WT1^+^ cells differentiation assessed by co-staining with vimentin. Representative images of WT1^+^/Vimentin^+^ cells at T0 and 48h of static and Tβ4-treated culture (D). Quantification of double positive cells in the epicardial layer (E), and in the whole slice (F). N of Pigs = 4-5, number of slices displayed in graphs. All graphs display data as mean with SEM. *=p ≤ 0.05, **=p ≤ 0.01. Scale bars, 100µm.

Tβ4 has been described as a potent stimulator of epicardial mobilization enabling differentiation in a variety of distinct phenotypes including smooth muscle cells, fibroblast and endothelial cells (*8*). To investigate this process in our *ex vivo* model, we quantified WT1^+^ cells co-expressing the mesenchymal marker vimentin. In freshly isolated slices, WT1^+^/Vimentin^+^ cells were relatively infrequent, and this pattern was maintained in untreated static culture (Fig. 5D). However, treatment with Tβ4 significantly increased the percentage of double positive cells both in the epicardial layer (T0 4.85±0.59, Stat 48h 1.72±0.62, Tβ4 18.67±6.39; Stat 48h vs. Tβ4 48h, p=0.0377) (Fig. 5E) and in the whole slice (Fig. 5F). To verify the bona fide epicardial origin of the WT1^+^ cells, we performed WT1/ Vimentin staining on Tβ4 treated myocardial slices after 48 hours of culture. Results showed an extremely limited presence of the WT1^+^ cells within the myocardial slices (Fig. S1)

Taken together, these data indicated a Tβ4-dependent activating effect on WT1^+^ epicardial cells, leading to an increase in migratory potential and mesenchymal differentiation. These results also supported the efficacy of this *ex vivo* model in enabling the study of WT1^+^ cell fate and their localization within the epicardium/myocardium interface.

## Discussion

The epicardium has emerged as one of the most relevant sources for a comprehensive and coordinated regeneration of cardiac tissue (*23*). Mouse studies demonstrated remarkable epicardial reactivation following apex amputation (*24*) and ischemia (*25*), but the reparative capacity of the epicardium is limited to early life and decreases dramatically in adulthood. In larger mammals, the lack of a representative model of heart remodeling limits the acquisition of knowledge which can be directly translated to improve patient care (*26*). New strategies are needed to stimulate the epicardial cell activation to unlock their regenerative capacity, therefore translating their promising experimental potential (*16*) into therapeutic improvements (*26*). Here, we describe a transformative organotypic 3D system representative of the epicardial/myocardial interface and containing all the cellular and extracellular components crucial to its physiology.

Myocardial slices, from both animal and human sources, have successfully enabled functional and pharmacological investigations into heart physiology (*19*–*21*). Our approach advances these existing models by including a fully functional epicardium, physiologically interfaced to the myocardium. To enable these preparations, we advanced existing embedding methods to protect the epicardial surface whilst achieving a flat cutting surface (*27, 28*). In fact, the epicardium has commonly been used to attach cardiac tissue blocks to specimen holders due to its flatness, which helps to align the tissue, preventing their damage during cutting (*29*), our methods achieves similar results, without losing the epicardium.

Epicardial slices maintain normal heart tissue architecture, as demonstrated by the preservation of functional markers and architecture in the myocardial portion of the slice, comparable with previous reports in myocardial slices (*19, 29, 30*). More importantly, slices present an intact epicardial monolayer expressing typical markers such as MSLN, E-cadherin and WT1. To our knowledge, this is the first comprehensive characterization of the epicardium in a large mammal heart. The majority of the epicardial cells on the surface of the slice expressed WT1. Although not entirely unique to the epicardial layer, this transcription factor has been extensively employed in lineage tracing studies in mice to identify epicardial progenitor cells (*31, 32*), and to described their plasticity after injury (*6, 33*). In mice, WT1^+^ cells account for over 90% of the epicardial cells in embryos, which is reduced to less than 25% in the adults (*6*). Comparison of our WT1^+^ results with other adult large mammal studies is hampered by the lack of available data, and we cannot therefore conclude to what extent epicardial activation is innate or may arise from the preparation process. In addition, epicardial cells and the immediate underlaying connective tissue strongly express MSLN, a surface marker characteristic of embryonic mesothelial progenitors originating fibroblasts and smooth muscle cells (*34*). Studies in mouse hearts indicated that MSLN expression is limited to the superficial epicardial layer in these animals (*6*). The combined widespread expression of WT1 and MSLN across the epicardium and the sub-epicardial layer suggests a sizable reservoir of epicardial progenitors in large adult mammal hearts.

We developed two separate protocols for the maintenance of epicardial slices in culture. The static culture system advances the well-established air-liquid interface approaches (*21*) by providing a pillared substrate as support to the myocardial face of the slice. The slices are pinned to pillars to maintain their shape and to maximize the contact surface area with the culture medium. We expect the tethering to increase slice viability since the application of static mechanical forces to myocardial slices is known to improve their performance in culture (*35*). The dynamic culture system perfuses the slices using a 3D printed insert that maintains a constant medium level, and a feedback loop control system that maintains controlled oxygen levels and stable pH in the culture.

Our results show that both culture systems preserve an unaltered cobblestone-shaped epicardial layer and sustain the overall slice viability and metabolic rate, comparable to previously reported myocardial slices from large mammals (*30, 36*). Notably, the more stable environment provided by the dynamic culture system promote proliferation of epicardial cells. Furthermore, dynamic culture preserves myocardial viability suggesting a its use to improve existing for myocardial slice protocols.

In order to investigate the suitability of the epicardial slices for pharmacological and mechanistic studies, we treated the slices with Tβ4 and monitored their response. Previous studies highlighted Tβ4 as potential therapeutic molecule to reduce myocardial damage after infarction, due to its ability to facilitate epicardial mobilization and promote neovascularization in the injured adult heart (*8, 22, 37*). In our *ex vivo* model, Tβ4 treatment preserves slice viability and increases epicardial cell proliferation, recapitulating previous data obtained in mice by intraperitoneal injection of Tβ4 after myocardial infarction (*22*). In addition, Tβ4 treatment induces a robust up-regulation of *WT1, Tbx18* and *TCF21* in the epicardial slices, while their expression remains low and mostly undetectable in the myocardial slices. These transcription factors are characteristic of epicardial cells during embryonic heart development (*6, 38*), and are re-expressed upon injury (*6*) and/or Tβ4 stimulation (*7, 39*), driving epicardial cell activation and contribution to cardiac repair (*7, 8, 37*). In mice, the reparative process was described to involve epicardial EMT and the subsequent invasion of the underlying myocardium, where the epicardial cells differentiated into various cardiac phenotypes (*7, 40*–*42*). Our model confirms the upregulation of the major epicardial EMT-inducible transcription factors, *Snai1, Snai2*, and *TWIST-1* (*15, 32*) in response to Tβ4 treatment. Concurrently, we detect a redistribution WT1^+^ cells towards the myocardium and an increased number of WT1^+^/vimentin^+^ cells following Tβ4 treatment. Importantly, the emergence of the WT1^+^/vimentin^+^ population was limited to the epicardial slices and did not occur in the myocardial slices, confirming the specific contribution of the epicardium. Overall, these results confirms in large animal heart tissues the pro-migratory and pro-differentiative effects of Tβ4 treatment in epicardial cells, as well as the reactivation of embryonic genes, that were previously described *in vitro* (*8*), and *in vivo* in mice (*6, 7, 10, 22, 43*). However, the increased number of mesenchymal WT1^+^/vimentin^+^ cells, seems to be in contrast with the antifibrotic effect attributed to Tβ4 (*11, 12*), and supports the need to further investigate the mechanisms of epicardial re-activation in large mammals. We cannot exclude that WT1^+^/vimentin^+^ cells define a population of epicardial-derived smooth muscle cells, although their position within the tissues do not indicate an association with blood vessels.

In conclusion, our study showed that living epicardial/myocardial slices can be obtained from porcine hearts and cultured effectively under different conditions. We also demonstrate that this epicardial/myocardial slice model can investigate epicardial cell activation and differentiation. The *ex vivo* model we developed preserves the native microenvironment of the epicardium and provides control of monitored and adjustable culture conditions, laying the path for the development of patient-relevant systems using human derived slices.

## Materials and Methods

### Experimental Design

This study aimed to develop an *ex vivo* 3D organotypic model of the epicardial/myocardial interface, which would enable studies directed at identifying mechanisms of adult epicardium reactivation. Our experiments verified the maintenance of the tissue architecture in the slices and the preservation of a living and healthy epicardial cells monolayer and myocardial tissue. Following Tβ4 stimulation, we evaluated the up-regulation of epicardial embryonic genes, and the EMT migration and the differentiation *in situ* of the WT1^+^ cells.

### Tissue samples

Swine hearts were obtained from The Pirbright Institute, UK. Animal experiments were carried out under the Home Office Animals (Scientific Procedures) Act (1986) (ASPA) and approved by the Animal Welfare and Ethical Review Board (AWERB) of The Pirbright Institute. The animals were housed in accordance with the Code of Practice for the Housing and Care of Animals Bred. Pigs of 4-6 weeks were euthanized by an overdose of 10ml pentobarbital (Dolethal 200mg/ml solution for injection, Vetoquinol UK Ltd). All procedures were conducted by Personal License holders who were trained and competent and under the Project License PPL70/8852. Hearts were rapidly excised and perfused with ice-cold cardioplegia solution (*44*) (NaCl 110mM; CaCl_2_ 1.2mM; KCl 16mM; MgCl_2_ 16mM; NaHCO_3_ 10mM; pH7.4).

### Slice preparation

Ventricle cubes of 8×8mm were embedded epicardium down in 5% liquified low melting agarose (ThermoScientific, TopVision Low Melting Point Agarose), dissolved in Normal Tyrode (NT) solution (NaCl 140mM; CaCl_2_ 1.8mM; KCl 6mM; MgCl_2_ 1mM; Glucose 10mM; HEPES 10mM; BDM 10mM; pH7.4).). To avoid epicardial cells damage embedding procedure was carried out on an ice-cold cushion of 2% agarose (Invitrogen UltraPure Agarose) dissolved in NT’s solution. Once cooled, embedded tissue cubes were mounted with the epicardium face up onto the specimen holder of a high precision vibratome (Leica, VT1200S) and cut to produce slices of 400-500µm thickness. After obtaining the epicardial slice, 400µm of tissue was discarded before cutting 2 to 6 myocardial slices of 300µm thickness. During cutting, the sample was constantly submerged in ice-cold and oxygenated (99.5% O_2_) NT solution. Blade advancement speed was set at 0.03mm/s and cutting amplitude at 1.50mm. After cutting, slices were incubated at least 30 minutes in room temperature (RT) NT solution before proceeding to culture or histological evaluation.

### Tissue culture

Slices were pinned using entomology pins (A1 - 0.14 × 10mm, Watkins & Doncaster), epicardium-up on 8mm-high Polydimethylsiloxane (PDMS) (SYLGARD 184) pillars cast at the bottom of a 100mm petri dish. Air-liquid interface was achieved by carefully adding culture medium (Medium 199 + 1X ITS Liquid Media Supplement + 1% Penicillin/Streptomycin Penicillin-Streptomycin + 10mM of BDM -all Sigma-Aldrich) to leave the epicardium exposed to the atmosphere. For epicardial cell re-activation experiments, culture medium was supplemented with 100 ng/ml of Tβ4 (Human Thymosin beta 4 peptide, Abcam). In the dynamic system, a custom 3D printed adaptor was inserted between the dish and the lid providing a connection to BioFlo 120 (Eppendorf) control station. Slices were cultured in an incubator with humidified air at 37 °C and 5% CO2.

### Live staining

Slices were incubated at room temperature with 10 µM Calcein AM cell-permeant dye (Invitrogen, Thermo Fisher Scientific) for 45 minutes under continuous shaking. Following washes, confocal images were collected from three random fields of each slice using a ×10 objective on a Nikon Eclipse Ti A1-A confocal laser scanning microscope. Z-stack confocal images were generated from picture taken at 5-10µm intervals, 1024×1024 pixels, from 7 to 30 sections. The percentage of area stained was measured on maximum intensity projection images using ImageJ.

### Slice morphology

Slices were fixed in 4% PFA (Paraformaldehyde, Santa Cruz Biotechnology) overnight (o/n) at 4°C, washed with phosphate buffer saline (PBS) and incubated overnight in 30% sucrose (Sigma-Aldrich) solution in PBS (w/v). Slices were frozen embedded in OCT Compound (Agar scientific) in dry ice and cryostat sectioned longitudinally obtaining 5µm thick sections.

Antigen retrieval was performed with microwave at 750 W for 15 min with citrate buffer (0.1M Citric Acid, pH 6.0) or water bath at 80°C for 30 min with tris-EDTA buffer (10mM Tris Base, 1mM EDTA Solution, pH 9.0) followed by permeabilization for nuclear antigens (0.1% Triton X-100 in PBS for 30 min) and blocking for 1 hour at room temperature with 20% Goat serum (Sigma-Aldrich) in PBS. Primary antibody incubation was performed overnight at 4°C (WT1 1:50, E-cadherin 1:50, CD31 1:100, NG2 1:500 (water bath antigen retrieval performed) all from Abcam; Mesothelin 1:100 from Novus Biologicals; α-Actinin (Sarcomeric) 1:800, α-SMA 1:400, PCNA 1:100 all from Sigma-Aldrich; Connexin 43 1:300, Vimentin 1:100 all from Thermo Fisher Scientific), followed by the appropriate Alexa Fluor (Thermo Fisher Scientific) secondary antibody diluted 1:200 for 1 hour at 37°C, and nuclei staining with DAPI (4′,6-diamidino-2-phenylindole, Merck) for 10 minutes at RT. Incubation with 0.1% Sudan Black (Sudan Black B, Santa Crus Biotechnology) in 70% ethanol (w/v) for 30 minutes at room temperature was performed to reduce tissue autofluorescence. Slides were then mounted in Fluoromount G (Invitrogen eBioscience Fluoromount G, Thermo Fisher Scientific) and imaged with Nikon Eclipse Ti A1-A confocal laser scanning microscope. Quantifications were performed on 3-7 random fields using ImageJ on 10x images.

### Gene expression analysis

RNA was extracted using Promega reliaprep RNA Miniprep System (Promega) from homogenate tissues, and reverse-transcribed using QuantiTect Reverse Transcription Kit (Qiagen). Real-time PCR was performed on QuantStudio 7 Flex Real-Time PCR System (Applied Biosystems) using QuantiTect SYBR Green PCR Kit (Qiagen) and the primers in Table 1 and results were normalized to the house-keeping gene β-2 microglobulin (B2M).

**Table 1.**
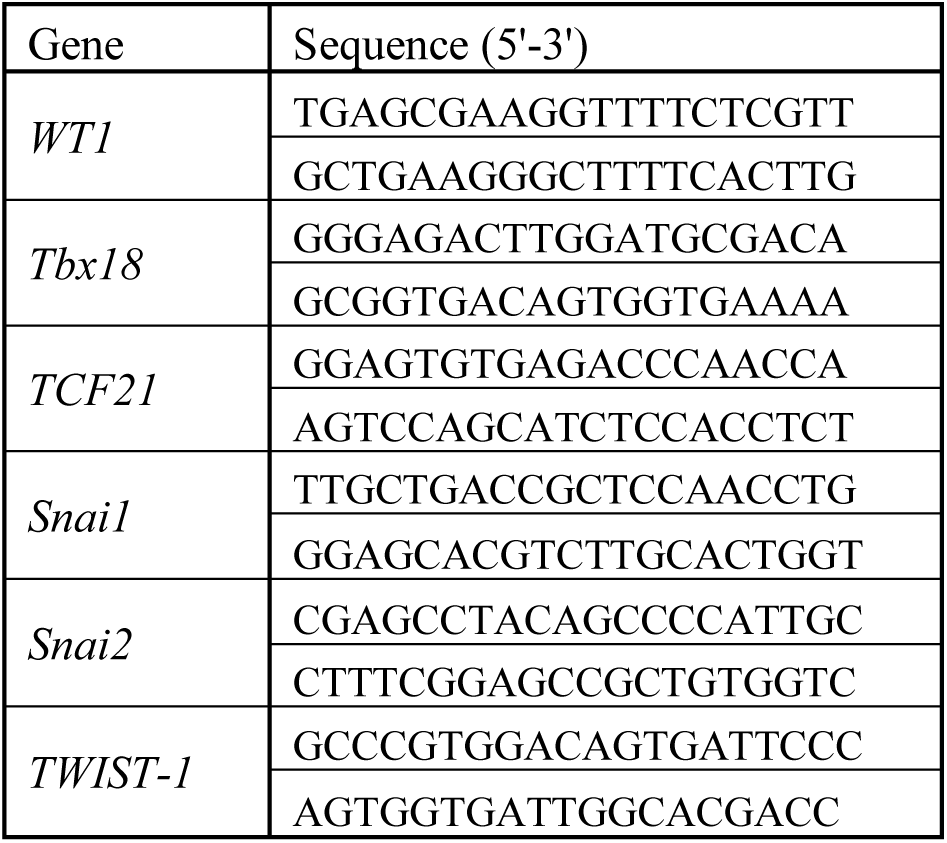
Lists of primers used in this study.

### Statistical analysis

Difference among groups were evaluated using one-way ANOVA or Kruskal–Wallis test, based on results from normality tests, followed by Fisher’s LSD post-hoc test (GraphPad Prism 8.1.2). A value of p<0.05 was considered as statistically significant. Data are presented as mean±SEM.

## Supplementary Materials

Fig. S1. Myocardial slices do not contain WT1^+^/vimentin^+^ cells after 48h of culture.

## Acknowledgments

**General**: We thank Drs L Dixton, A Reis and M Henstock from the Pirbright Institute (Pirbright, UK) and the personnel at Newman Abattoir (Farnborough, UK) for their support in procuring the animal tissues. Prof John McVey and the Department of Biochemical Sciences at the University of Surrey, especially the technical team, for their continuing support.

## Author contributions

DM conceived and planned the experiments and analyzed the data. RSM and RDJ provided key experimental expertise. DM and PC2 wrote the manuscript and prepared the figures with support of CC. PC2, PC1 and CC conceived the original idea and supervised the project.

## Data and materials availability

TBD.

## Supplementary Materials

**Fig. S1.**
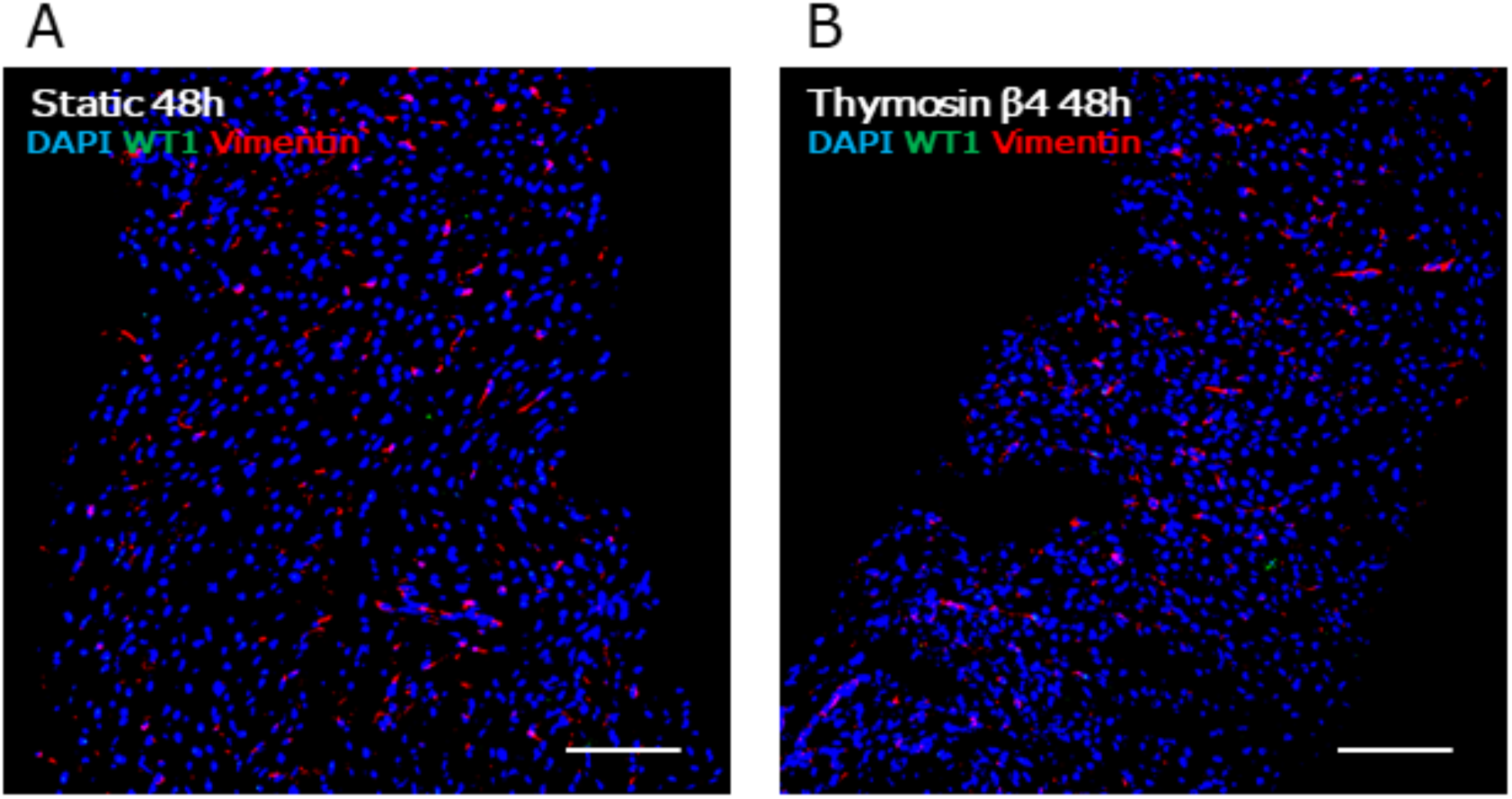
Myocardial slices do not contain WT1^+^/vimentin^+^ cells after 48h of culture. Representative images of WT1/Vimentin double-staining at 48h of static (A) and Tβ4-treated culture of myocardial slices (B). WT1 staining is undetectable in the tissue. Scale bars, 100µm.

## References

1. J. M. Pérez-Pomares, D. Macías, L. García-Garrido, R. Muñoz-Chápuli, The origin of the subepicardial mesenchyme in the avian embryo: An immunohistochemical and quail-chick chimera study. Dev. Biol. (1998), doi:10.1006/dbio.1998.8949.

2. S. R. Ali, S. Ranjbarvaziri, M. Talkhabi, P. Zhao, A. Subat, A. Hojjat, P. Kamran, A. M. S. Müller, K. S. Volz, Z. Tang, K. Red-Horse, R. Ardehali, Developmental heterogeneity of cardiac fibroblasts does not predict pathological proliferation and activation. Circ. Res. 115, 625–635 (2014).

3. Y. Yamaguchi, S. Cavallero, M. Patterson, H. Shen, J. Xu, S. R. Kumar, H. M. Sucov, Adipogenesis and epicardial adipose tissue: A novel fate of the epicardium induced by mesenchymal transformation and PPARγ activation. Proc. Natl. Acad. Sci. U. S. A. 112, 2070–2075 (2015).

4. K. S. Volz, A. H. Jacobs, H. I. Chen, A. Poduri, A. S. McKay, D. P. Riordan, N. Kofler, J. Kitajewski, I. Weissman, K. Red-Horse, Pericytes are progenitors for coronary artery smooth muscle. Elife. 4, 1–22 (2015).

5. B. van Wijk, Q. D. Gunst, A. F. M. Moorman, M. J. B. van den Hoff, Cardiac Regeneration from Activated Epicardium. PLoS One. 7 (2012), doi:10.1371/journal.pone.0044692.

6. B. Zhou, L. B. Honor, H. He, M. Qing, J. H. Oh, C. Butterfield, R. Z. Lin, J. M. Melero-Martin, E. Dolmatova, H. S. Duffy, A. Von Gise, P. Zhou, Y. W. Hu, G. Wang, B. Zhang, L. Wang, J. L. Hall, M. A. Moses, F. X. McGowan, W. T. Pu, Adult mouse epicardium modulates myocardial injury by secreting paracrine factors. J. Clin. Invest. (2011), doi:10.1172/JCI45529.

7. N. Smart, S. Bollini, K. N. Dubé, J. M. Vieira, B. Zhou, S. Davidson, D. Yellon, J. Riegler, A. N. Price, M. F. Lythgoe, W. T. Pu, P. R. Riley, De novo cardiomyocytes from within the activated adult heart after injury. Nature. 474, 640–644 (2011).

8. N. Smart, C. A. Risebro, A. A. D. Melville, K. Moses, R. J. Schwartz, K. R. Chien, P. R. Riley, Thymosin β4 induces adult epicardial progenitor mobilization and neovascularization. Nature. 445, 177–182 (2007).

9. J. M. Vieira, S. Howard, C. Villa del Campo, S. Bollini, K. N. Dubé, M. Masters, D. N. Barnette, M. Rohling, X. Sun, L. E. Hankins, D. Gavriouchkina, R. Williams, D. Metzger, P. Chambon, T. Sauka-Spengler, B. Davies, P. R. Riley, BRG1-SWI/SNF-dependent regulation of the Wt1 transcriptional landscape mediates epicardial activity during heart development and disease. Nat. Commun. 8, 16034 (2017).

10. N. Smart, S. Bollini, K. N. Dubé, J. M. Vieira, B. Zhou, J. Riegler, A. N. Price, M. F. Lythgoe, S. Davidson, D. Yellon, W. T. Pu, P. R. Riley, Myocardial regeneration: Expanding the repertoire of thymosin β4 in the ischemic heart. Ann. N. Y. Acad. Sci. 1269, 92–101 (2012).

11. S. Kumar, S. Gupta, Thymosin beta 4 prevents oxidative stress by targeting antioxidant and anti-apoptotic genes in cardiac fibroblasts. PLoS One (2011), doi:10.1371/journal.pone.0026912.

12. M. A. Evans, N. Smart, K. N. Dubé, S. Bollini, J. E. Clark, H. G. Evans, L. S. Taams, R. Richardson, M. Lévesque, P. Martin, K. Mills, J. Riegler, A. N. Price, M. F. Lythgoe, P. R. Riley, Thymosin β4-sulfoxide attenuates inflammatory cell infiltration and promotes cardiac wound healing. Nat. Commun. (2013), doi:10.1038/ncomms3081.

13. J. M. González-Rosa, C. E. Burns, C. G. Burns, Zebrafish heart regeneration: 15 years of discoveries. Regeneration. 4, 105–123 (2017).

14. B. J. Haubner, J. Schneider, U. Schweigmann, T. Schuetz, W. Dichtl, C. Velik-Salchner, J. I. Stein, J. M. Penninger, Functional Recovery of a Human Neonatal Heart after Severe Myocardial Infarction. Circ. Res. 118, 216–221 (2016).

15. O. M. Martínez-Estrada, L. A. Lettice, A. Essafi, J. A. Guadix, J. Slight, V. Velecela, E. Hall, J. Reichmann, P. S. Devenney, P. Hohenstein, N. Hosen, R. E. Hill, R. Mũoz-Chapuli, N. D. Hastie, Wt1 is required for cardiovascular progenitor cell formation through transcriptional control of Snail and E-cadherin. Nat. Genet. (2010), doi:10.1038/ng.494.

16. A. T. Moerkamp, K. Lodder, T. Van Herwaarden, E. Dronkers, C. K. E. Dingenouts, F. C. Tengström, T. J. Van Brakel, M. J. Goumans, A. M. Smits, Human fetal and adult epicardial-derived cells: A novel model to study their activation. Stem Cell Res. Ther. (2016), doi:10.1186/s13287-016-0434-9.

17. W. C. Claycomb, Biochemical aspects of cardiac muscle differentiation. Possible control of deoxyribonucleic acid synthesis and cell differentiation by adrenergic innervation and cyclic adenosine 3’:5’ monophosphate. J. Biol. Chem. (1976).

18. T. P. de Boer, P. Camelliti, U. Ravens, P. Kohl, Myocardial tissue slices: Organotypic pseudo-2D models for cardiac research & development. Future Cardiol. (2009), doi:10.2217/fca.09.32.

19. P. Camelliti, S. A. Al-Saud, R. T. Smolenski, S. Al-Ayoubi, A. Bussek, E. Wettwer, N. R. Banner, C. T. Bowles, M. H. Yacoub, C. M. Terracciano, Adult human heart slices are a multicellular system suitable for electrophysiological and pharmacological studies. J. Mol. Cell. Cardiol. 51, 390–398 (2011).

20. A. Bussek, E. Wettwer, T. Christ, H. Lohmann, P. Camelliti, U. Ravens, Tissue slices from adult mammalian hearts as a model for pharmacological drug testing. Cell. Physiol. Biochem. 24, 527–536 (2009).

21. M. Brandenburger, J. Wenzel, R. Bogdan, D. Richardt, F. Nguemo, M. Reppel, H. Terlau, A. Dendorfer, Organotypic slice culture from human adult ventricular myocardium, 50–59 (2012).

22. N. Smart, C. A. Risebro, J. E. Clark, E. Ehler, L. Miquerol, A. Rossdeutsch, M. S. Marber, P. R. Riley, Thymosin β4 facilitates epicardial neovascularization of the injured adult heart. Ann. N. Y. Acad. Sci. 1194, 97–104 (2010).

23. A. Wessels, J. M. Pérez-Pomares, The Epicardium and Epicardially Derived Cells (EPDCs) as Cardiac Stem Cells. Anat. Rec. - Part A Discov. Mol. Cell. Evol. Biol. (2004), doi:10.1002/ar.a.10129.

24. E. R. Porrello, A. I. Mahmoud, E. Simpson, J. A. Hill, J. A. Richardson, E. N. Olson, H. A. Sadek, Transient regenerative potential of the neonatal mouse heart. Science (80-.). (2011), doi:10.1126/science.1200708.

25. E. R. Porrello, A. I. Mahmoud, E. Simpson, B. A. Johnson, D. Grinsfelder, D. Canseco, P. P. Mammen, B. A. Rothermel, E. N. Olson, H. A. Sadek, Regulation of neonatal and adult mammalian heart regeneration by the miR-15 family. Proc. Natl. Acad. Sci. U. S. A. (2013), doi:10.1073/pnas.1208863110.

26. T. R. Heallen, Z. A. Kadow, J. H. Kim, J. Wang, J. F. Martin, Stimulating Cardiogenesis as a Treatment for Heart Failure. Circ. Res. (2019), doi:10.1161/CIRCRESAHA.118.313573.

27. Q. Wen, K. Gandhi, R. A. Capel, G. Hao, C. O’Shea, G. Neagu, S. Pearcey, D. Pavlovic, D. A. Terrar, J. Wu, G. Faggian, P. Camelliti, M. Lei, Transverse cardiac slicing and optical imaging for analysis of transmural gradients in membrane potential and Ca2+ transients in murine heart. J. Physiol. (2018), doi:10.1113/JP276239.

28. M. Halbach, F. Pillekamp, K. Brockmeier, J. Hescheler, J. Müller-Ehmsen, M. Reppel, Ventricular slices of adult mouse hearts - A new multicellular in vitro model for electrophysiological studies. Cell. Physiol. Biochem. (2006), doi:10.1159/000095132.

29. S. A. Watson, M. Scigliano, I. Bardi, R. Ascione, C. M. Terracciano, F. Perbellini, Preparation of viable adult ventricular myocardial slices from large and small mammals. Nat. Protoc. 12, 2623–2639 (2017).

30. Q. Ou, Z. Jacobson, R. R. E. Abouleisa, X. L. Tang, S. M. Hindi, A. Kumar, K. N. Ivey, G. Giridharan, A. El-Baz, K. Brittian, B. Rood, Y. H. Lin, S. A. Watson, F. Perbellini, T. A. McKinsey, B. G. Hill, S. P. Jones, C. M. Terracciano, R. Bolli, T. M. A. Mohamed, Physiological Biomimetic Culture System for Pig and Human Heart Slices. Circ. Res. (2019), doi:10.1161/CIRCRESAHA.119.314996.

31. B. Zhou, Q. Ma, S. Rajagopal, S. M. Wu, I. Domian, J. Rivera-Feliciano, D. Jiang, A. Von Gise, S. Ikeda, K. R. Chien, W. T. Pu, Epicardial progenitors contribute to the cardiomyocyte lineage in the developing heart. Nature (2008), doi:10.1038/nature07060.

32. B. Zhou, A. von Gise, Q. Ma, Y. W. Hu, W. T. Pu, Genetic fate mapping demonstrates contribution of epicardium-derived cells to the annulus fibrosis of the mammalian heart. Dev. Biol. (2010), doi:10.1016/j.ydbio.2009.12.007.

33. B. Zhou, W. T. Pu, Genetic Cre-loxP assessment of epicardial cell fate using Wt1-Driven cre alleles. Circ. Res. (2012), doi:10.1161/CIRCRESAHA.112.275784.

34. Y. Rinkevich, T. Mori, D. Sahoo, P. X. Xu, J. R. Bermingham, I. L. Weissman, Identification and prospective isolation of a mesothelial precursor lineage giving rise to smooth muscle cells and fibroblasts for mammalian internal organs, and their vasculature. Nat. Cell Biol. (2012), doi:10.1038/ncb2610.

35. S. A. Watson, J. Duff, I. Bardi, M. Zabielska, S. S. Atanur, R. J. Jabbour, A. Simon, A. Tomas, R. T. Smolenski, S. E. Harding, F. Perbellini, C. M. Terracciano, Biomimetic electromechanical stimulation to maintain adult myocardial slices in vitro. Nat. Commun. (2019), doi:10.1038/s41467-019-10175-3.

36. F. Perbellini, S. A. Watson, M. Scigliano, S. Alayoubi, S. Tkach, I. Bardi, N. Quaife, C. Kane, N. P. Dufton, A. Simon, M. B. Sikkel, G. Faggian, A. M. Randi, J. Gorelik, S. E. Harding, C. M. Terracciano, Investigation of cardiac fibroblasts using myocardial slices. Cardiovasc. Res. (2018), doi:10.1093/cvr/cvx152.

37. I. Bock-Marquette, A. Saxena, M. D. White, J. M. DiMaio, D. Srivastava, Thymosin β4 activates integrin-linked kinase and promotes cardiac cell migration, survival and cardiac repair. Nature. 432, 466–472 (2004).

38. A. Acharya, S. T. Baek, G. Huang, B. Eskiocak, S. Goetsch, C. Y. Sung, S. Banfi, M. F. Sauer, G. S. Olsen, J. S. Duffield, E. N. Olson, M. D. Tallquist, The bHLH transcription factor Tcf21 is required for lineagespecific EMT of cardiac fibroblast progenitors. Dev. 139, 2139–2149 (2012).

39. S. Chen, M. Shimoda, J. Chen, P. A. Grayburn, Stimulation of adult resident cardiac progenitor cells by durable myocardial expression of thymosin beta 4 with ultrasound-targeted microbubble delivery. Gene Ther. 20, 225–233 (2013).

40. A. von Gise, B. Zhou, L. B. Honor, Q. Ma, A. Petryk, W. T. Pu, WT1 regulates epicardial epithelial to mesenchymal transition through β-catenin and retinoic acid signaling pathways. Dev. Biol. (2011), doi:10.1016/j.ydbio.2011.05.668.

41. M. Zamora, J. Männer, P. Ruiz-Lozano, Epicardium-derived progenitor cells require β-catenin for coronary artery formation. Proc. Natl. Acad. Sci. U. S. A. (2007), doi:10.1073/pnas.0702415104.

42. N. Smart, C. A. Risebro, A. A. D. Melville, K. Moses, R. J. Schwartz, K. R. Chien, P. R. Riley, Thymosin β-4 is essential for coronary vessel development and promotes neovascularization via adult epicardium. Ann. N. Y. Acad. Sci. 1112, 171–188 (2007).

43. B. Zhou, L. B. Honor, Q. Ma, J. H. Oh, R. Z. Lin, J. M. Melero-Martin, A. von Gise, P. Zhou, T. Hu, L. He, K. H. Wu, H. Zhang, Y. Zhang, W. T. Pu, Thymosin beta 4 treatment after myocardial infarction does not reprogram epicardial cells into cardiomyocytes. J. Mol. Cell. Cardiol. 52, 43–47 (2012).

44. C. Kang, Y. Qiao, G. Li, K. Baechle, P. Camelliti, S. Rentschler, I. R. Efimov, Human Organotypic Cultured Cardiac Slices: New Platform For High Throughput Preclinical Human Trials. Sci. Rep. (2016), doi:10.1038/srep28798.

